# Conserved role of the SERK–BIR module in development and immunity across land plants

**DOI:** 10.1101/2024.08.14.607940

**Authors:** Yijia Yan, Jaqueline Mellüh, Martin A. Mecchia, Hyung-Woo Jeon, Katharina Melkonian, Clemens Holzberger, Anne Harzen, Sara Christina Stolze, Rainer Franzen, Yuki Hirakawa, Ana I. Caño Delgado, Hirofumi Nakagami

## Abstract

SOMATIC EMBRYOGENESIS RECEPTOR-LIKE KINASES (SERKs), which are subfamily II of leucine-rich repeat receptor-like kinases (LRR-RLKs), play diverse roles in development and immunity in the angiosperm *Arabidopsis thaliana*. AtSERKs act as co-receptors for many LRR-RLKs, including BRASSINOSTEROID INSENSITIVE 1 (BRI1) and FLAGELLIN SENSITIVE 2 (FLS2).^1–4^ The conserved tyrosine (Y) residue in AtSERK3 is crucial for signaling specificity in differentiating BRI1- and FLS2-mediated pathways.^5^ BRI1-ASSOCIATED RECEPTOR KINASE 1 (BAK1)-INTERACTING RECEPTOR-LIKE KINASES (BIRs) interact with SERKs under resting conditions, negatively regulating SERK-mediated pathways.^6,7^ SERK and BIR are highly conserved in land plants, whereas BRI1 and FLS2 homologs are absent or poorly conserved in bryophyte lineages.^8,9^ The biological functions of SERK homologs in non-flowering plants are largely unknown. The genome of the liverwort *Marchantia polymorpha* encodes single homologs for SERK and BIR, namely MpSERK and MpBIR.^9^ We here show that Mp*serk* disruptants display growth and developmental defects with no observable sexual or vegetative reproduction. Complementation analysis revealed a contribution of the conserved Y residue of MpSERK to growth. Proximity labelling-based interactomics identified MpBIR as a MpSERK interactor. Mp*bir* disruptants displayed defects in reproductive organ development. Patterns of development- and immunity-related gene expression in Mp*serk* and Mp*bir* were antagonistic, suggesting that MpBIR functions as a MpSERK repressor. The pathogenic bacterium *Pseudomonas syringae* pv. *tomato* DC3000 grew poorly on Mp*bir*, indicating a significant role of the MpSERK1MpBIR module in immunity. Taken together, we propose that the SERK–BIR functional module was already regulating both development and immunity in the last common ancestor of land plants.

## Results and discussion

### MpSERK is required for thalli development and reproduction

The *Marchantia polymorpha* genome encodes three LRR-RLK subfamily II (LRR-RLK II) members. Among the three LRR-RLK IIs, only Mp7g09160 is orthologous to *Arabidopsis thaliana* SERKs, and therefore Mp7g09160 has been named MpSERK.^10^ The two other members are orthologous to AtCIKs and AtAPEX, respectively, and MpCIK is likely to function as a co-receptor of MpCLV1.^10–12^ The LRR-RLK IIs found in the transcriptome of the Zygnematophyceae alga *Spirogyra pratensis* are orthologous to all of the SERK, CIK, and APEX members, although it can be most highly related to SERK.^10,13^ It is very likely that the most recent common ancestor (MRCA) of land plants had three LRR-RLK IIs resulting from gene duplications (Figure 1A).

**Figure 1.**
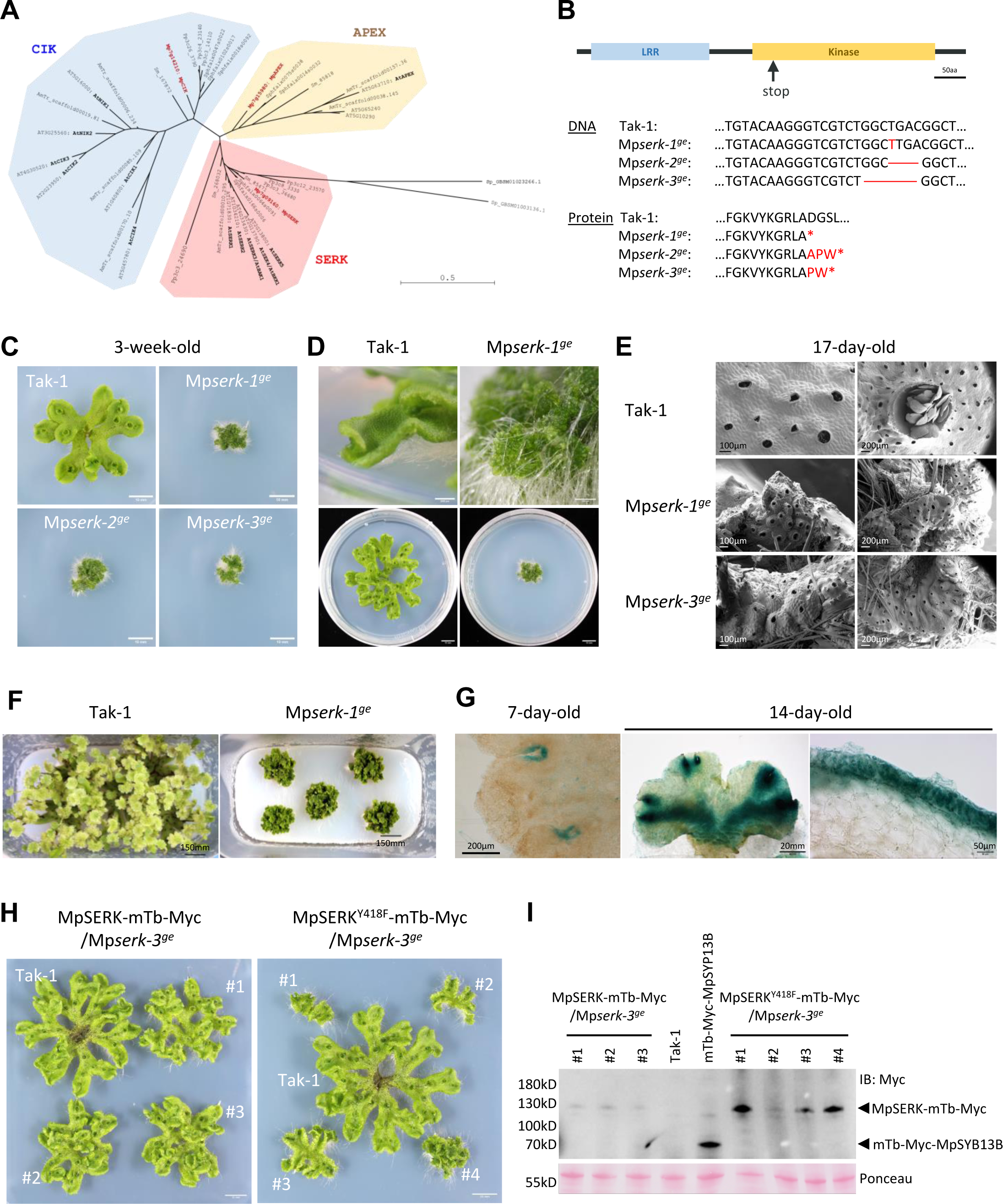
Mp*SERK* is required for thalli development and reproduction. (A) Unrooted phylogenetic tree of the subfamily II LRR-RLKs in plants. The amino acid sequences of the kinase domain were used for analysis. (B) Schematic representation of Mp*SERK* disruptions in the Mp*serk-1^ge^,* Mp*serk-2^ge^*, and Mp*serk-3^ge^*. Premature translation termination in the kinase domain of MpSERK is indicated by an arrow. (C) Three-week-old Tak-1, Mp*serk-1^ge^,* Mp*serk-2^ge^*, and Mp*serk-3^ge^* grown on agar plates. Thalli were grown from single gemma (Tak-1) or small fragments of thalli (Mp*serk^ge^*mutants). Scale bars, 10 mm. (D) Four-week-old Tak-1 and Mp*serk-1^ge^*grown on agar plates. Scale bars, 2000 μm (upper left panel), 1000 μm (upper right panel), 10 mm (lower panels). (E) Scanning electronic microscope (SEM) images of surfaces of 17-day-old Tak-1, Mp*serk-1^ge^*, and Mp*serk-3^ge^*. (F) Induction of gametangiophores in Tak-1 and Mp*serk-1^ge^*. All plants were 38 days old. Thalli were grown from single gemma (Tak-1) or small fragments of thalli (Mp*serk-1^ge^*) under constant white light supplemented with far-red light. (G) GUS staining images of thalli harboring *_pro_*Mp*SERK:GUS*. Right image shows the cross-sectional view of 14-day-old thallus. (H) Thirty-one-day-old Tak-1, *_pro_*Mp*SERK:*Mp*SERK*-miniTurbo-Myc/Mp*serk-3^ge^*, and *_pro_*Mp*SERK:*Mp*SERK^Y418F^*-miniTurbo-Myc/Mp*serk-3^ge^* grown on agar plates. Thalli were grown from single gemma. Scale bars, 10 mm. (I) Protein expression levels of MpSERK-miniTurbo-Myc or MpSERK^Y418F^-miniTurbo-Myc in the transgenic plants shown in (H). Myc-tagged proteins were detected using anti-Myc antibody and indicated by arrows. Ponceau S-stained membrane is shown as a loading control.

To examine functions of the single SERK in *M. polymorpha*, we generated CRISPR/Cas9-based MpSERK loss-of-function mutants in the *M. polymorpha* Tak-1 background. We obtained independent mutant alleles that lack a large part of the intracellular kinase domain (Figure1B). All the Mp*serk^ge^*mutants displayed very similar severe growth and developmental defects, which could be restored by expression of MpSERK fused to miniTurbo (mTb) biotin ligase and Myc-tag at its C-terminus (MpSERK-mTb-Myc) under its own promoter (Figures 1C and 1H). Thalli of Mp*serk^ge^* mutants showed enhanced branching and wavy surfaces compared to wild-type Tak-1 (Figures 1C11E). The Mp*serk^ge^* mutants were capable of developing air chambers and smooth rhizoids, while failing to develop gemma cups (Figures 1C11E). Far-red light irradiation did not induce gametangiophore formation in Mp*serk^ge^* mutants after up to 38 days (Figure 1F). These results indicate that MpSERK plays a role in initiating vegetative and sexual reproduction. GUS reporter-based promoter analysis indicated that MpSERK is primarily expressed in meristematic regions, consistent with the developmental defects of Mp*serk^ge^* mutants likely being caused by mis-regulations in this area.^14–17^ Expression of MpSERK was also observed in assimilatory filaments, which may imply a role in *M. polymorpha* immunity as suggested by a recent study (Figure 1G).^18^

*Arabidopsis thaliana* SERK3/BAK1 (AtSERK3/AtBAK1) functions as a co-receptor of pattern-recognition receptors (PRRs), many of which are LRR-RLK XII members such as AtFLS2, and contributes to defense against pathogenic bacteria.^4^ Because of its severe developmental phenotype, it was not feasible to appropriately compare the growth of the pathogenic bacterium *Pseudomonas syringae* pv. *tomato* DC3000 (*Pto* DC3000) on Mp*serk^ge^* thalli with growth on Tak-1.^18,19^ In *A. thaliana*, phosphorylation of AtBAK1 at Y403 is required for immune signaling but not for brassinosteroid signaling.^5^ This Y residue, which is located at the kinase domain VI, is highly conserved in SERK homologs including MpSERK.^5^ Therefore, we asked whether the growth and developmental defects of Mp*serk^ge^*could be uncoupled from defects in immune signaling by mutating the Y residue of MpSERK. Contrary to our expectations, expression of MpSERK^Y418F^-mTb-Myc under its own promoter could not complement the growth and developmental phenotypes to the same extent as MpSERK-mTb-Myc (Figure 1H). Protein expression of MpSERK^Y418F^-mTb-Myc could be confirmed, which was even higher compared to MpSERK-mTb-Myc (Figure 1I). Therefore, in *M. polymorpha*, the conserved Y residue appears to play a role in signaling related to growth but not immunity. It is also possible that Y418 regulates MpSERK protein stability and that the observed phenotype was caused by overaccumulation of MpSERK. In any case, the importance of the conserved Y residue for the molecular functions of SERK seems to be evolutionarily conserved, and other approaches need to be taken to investigate a possible contribution of MpSERK to immunity.

### MpSERK interacts with MpBIR

Assuming that MpSERK functions as a co-receptor of LRR-RLKs in *M. polymorpha*, we performed miniTurbo-based interactomics to identify LRR-RLKs that may interact with MpSERK or MpSERKY^418F^. Tak-1 and plants expressing plasma membrane-localized mTb-Myc-MpSYP13B were used as controls.^20^ When we used Tak-1 as a control, we could identify 16 and 15 LRR-RLKs that potentially interact or form a complex with MpSERK and MpSERKY^418F^, respectively (Figures 2A and 2B; Data S1A and S1B). We did not identify any LRR-RLK XII members, probably because the plants were not exposed to bacterial elicitors that can be recognized by PRRs and induce receptor complex formation. It should be noted that peptidoglycan is thus far the only bacterial component described to trigger immune responses in *M. polymorpha*.^18^ Further comparison to the interactome of MpSYP13B suggested that MpBIR could be the single very specific LRR-RLK that interacts with MpSERK in the resting state (Figure S1; Data S1D and S1E). Interestingly, MpTDR and MpRGI1 were identified as potential interactors of MpSERKY^418F^ but not of MpSERK (Figures 2C and S1; Data S1C–S1E).^21,22^ The Y418 of MpSERK may contribute to MpTDR- and MpRGI1-mediated signaling, and the observed growth phenotype of MpSERKY^418F^/Mp*serk^ge^* plants might be due to mis-regulation of MpTDR and MpRGI1 pathways.

**Figure 2.**
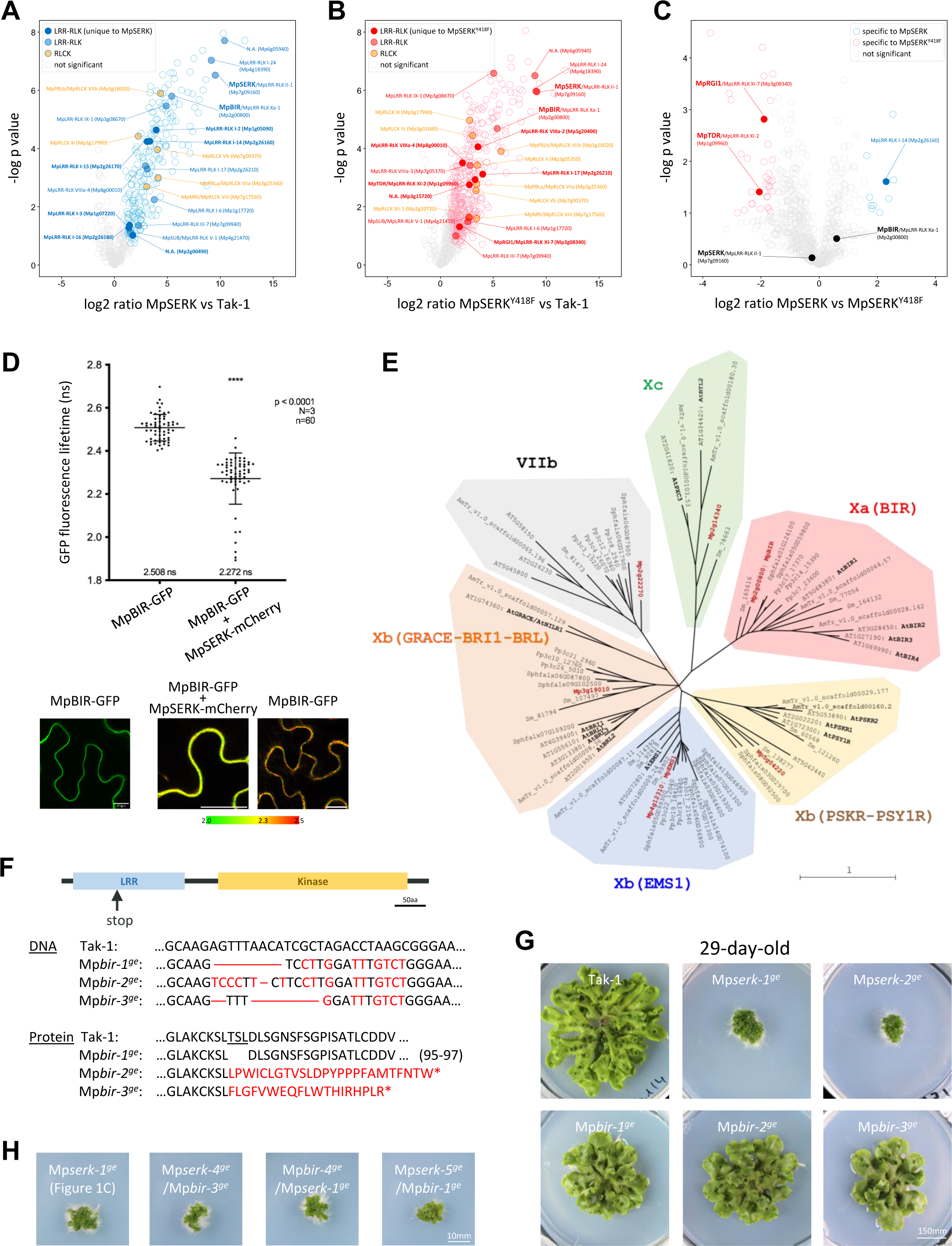
MpSERK interacts with MpBIR. (A–C) Interactome analysis of MpSERK and MpSERK^Y418F^. (A) Proteins significantly enriched using MpSERK as a bait are indicated in blue circles. LRR-RLKs and RLCKs are highlighted as filled circles with blue and orange, respectively. See Data S1A. (B) Proteins significantly enriched using MpSERK^Y418F^ as a bait are indicated in red circles. LRR-RLKs and RLCKs are highlighted as filled circles with red and orange, respectively. See Data S1B. (C) Proteins significantly enriched with MpSERK and MpSERK^Y418F^ are indicated in blue and red, respectively, when comparing between interactomes of MpSERK and MpSERK^Y418F^. See Data S1C. See also Figure S1, Data S1D, and Data S1E. (D) Interaction of MpBIR-GFP and MpSERK-mCherry in *N. benthamiana* leaves. Upper graph shows mean fluorescence lifetime (τ, ns) of MpBIR-GFP when expressed alone or along with MpSERK-mCherry. Significant difference calculated by one-way ANOVA followed by Dunnett’s test is indicated with asterisks (****, p < 0.0001). Error bars, standard deviation; n, number of measurements; N, number of independent experiments. In a lower left panel, a representative confocal image of *N. benthamiana* leaf cells expressing MpBIR-GFP is shown. Green pseudo-color indicates the fluorescence from GFP. Scale bar, 20 μm. In lower mid and right panels, GFP lifetimes are shown using pseudo-color according to the color code ranging from 2.0 ns (Green) to 2.5 ns (red). The respective lifetime values measured for MpBIR-GFP expressed alone or co-expressed with MpSERK-mCherry are indicated based on the color scales. Scale bars, 25 μm. (E) Unrooted phylogenetic tree of LRR-RLKs from the subfamily X and VI in plants. The amino acid sequences of the kinase domain were used for analysis. (F) Schematic representation of Mp*BIR* disruption in the Mp*bir-1^ge^*, Mp*bir-2^ge^*, and Mp*bir-3^ge^*. Premature translation termination in the LRR domain of MpBIR is indicated by an arrow. (G) Twenty-nine-day-old Tak-1, Mp*bir-1^ge^*, Mp*bir-2^ge^*, Mp*bir-3^ge^*, Mp*serk-1^ge^*, and Mp*serk-2^ge^* grown on agar plates. Thalli were grown from single gemma (Tak-1 and Mp*bir^ge^* mutants) or small fragments of thalli (Mp*serk^ge^* mutants). (H) Three-week-old Tak-1, Mp*serk-1^ge^*, and Mp*serk^ge^/bir^ge^* double mutants grown on agar plates. Thalli were grown from single gemma (Tak-1) or small fragments of thalli (Mp*serk-1^ge^* and Mp*serk^ge^/bir^ge^* double mutants). The image used here for Mp*serk-1^ge^* is the same image shown in Figure 1C as indicated in the figure. See also Figure S2.

To verify the interaction of MpSERK and MpBIR, fusion proteins fluorescently tagged at their C-terminus were transiently expressed in *Nicotiana benthamiana* leaves using *Agrobacterium*-mediated transient transformation. Förster resonance energy transfer (FRET) and fluorescence lifetime imaging microscopy (FLIM) confirmed that MpSERK and MpBIR interact in plants at cell surfaces (Figure 2D). These results indicate that MpSERK and MpBIR function as a module. The *M. polymorpha* genome encodes a single BIR homolog, while BIR homologs or LRR-RLK Xa members are so far found only in the genomes of land plants (Figure 2E). BIR homologs from gymnosperms, ferns, lycophytes, and bryophytes are all closely related to AtBIR1.^8^ It is thus likely that the SERK1BIR module was already established in the MRCA of land plants.

To investigate functions of the single BIR and its relationship with MpSERK in *M. polymorpha*, we have generated CRISPR/Cas9-based MpBIR loss-of-function mutants, Mp*bir^ge^*, and Mp*serk^ge^*/Mp*bir^ge^* double mutants (Figures 2F–2H). Three obtained Mp*bir^ge^* mutants displayed very similar, rather mild growth and developmental defects (Figures 2F and 2G). Interestingly, one of the mutant alleles, Mp*bir-1^ge^*, had only a three-amino acid deletion at the extracellular LRR domain of MpBIR (Figure 2F). This small deletion may affect proper MpSERK1MpBIR complex formation dynamics.^7^ All Mp*serk^ge^*/Mp*bir^ge^* double mutants phenocopied Mp*serk^ge^*, showing that Mp*serk^ge^* is epistatic to Mp*bir^ge^* (Figure 2H). Moreover, overaccumulation of MpBIR in Tak-1 resulted in the Mp*serk^ge^*-like phenotype (Figure S2). Taken together, a major molecular function of MpBIR could be the suppression of MpSERK activity by direct physical interactions, as in the case of *A. thaliana* SERK1BIR modules.^7^

### MpBIR negatively regulates immunity and is required for gemma cup and gametangiophore development

Given that BIRs generally function as negative regulators of SERKs in *A. thaliana*, we hypothesized that MpSERK-mediated pathways could be activated in Mp*bir^ge^* mutants in the absence of pathogen challenge. Therefore, we profiled transcriptomes of Tak-1, Mp*serk^ge^*, and Mp*bir^ge^* grown on agar plates under our normal growth conditions to investigate any potential divergence in gene expression between Mp*serk^ge^* and Mp*bir^ge^*. Strikingly, 40% of the differentially expressed genes in Mp*serk^ge^* and Mp*bir^ge^* showed antagonistic expression patterns compared to Tak-1, supporting our hypothesis (Figure 3A: Cluster 1, 3, and 10; Data S1F). Genes in clusters 1 and 3 were particularly interesting because these were induced in Mp*bir^ge^* and reduced in Mp*serk^ge^* compared to Tak-1 (Figure 3A). Gene ontology (GO) analysis showed a significant enrichment of growth- and development-related GO terms in cluster 3, supporting the observed roles of MpSERK in growth and development (Figure 3B; Data S1G). Defense-related GO terms were significantly enriched in cluster 1 (Figure 3B; Data S1H). In cluster 1, there was a slight difference in gene expression levels between Tak-1 and Mp*serk^ge^* compared to cluster 3 (Figure 3A). This is not surprising, because the analyzed Tak-1 plants were grown in the absence of immune activation. Plant cell-derived ligands that regulate growth and development are likely to activate MpSERK-dependent pathways under non-stimulating conditions, and therefore a clear difference between Tak-1 and Mp*serk^ge^* could be observed in cluster 3. These results suggest that MpBIR plays a role in restricting undesired MpSERK activation when ligands or stimuli are absent. MpBIR negatively regulates defense-related gene expression possibly through repression of MpSERK. In other words, MpSERK may positively contribute to immunity.

**Figure 3.**
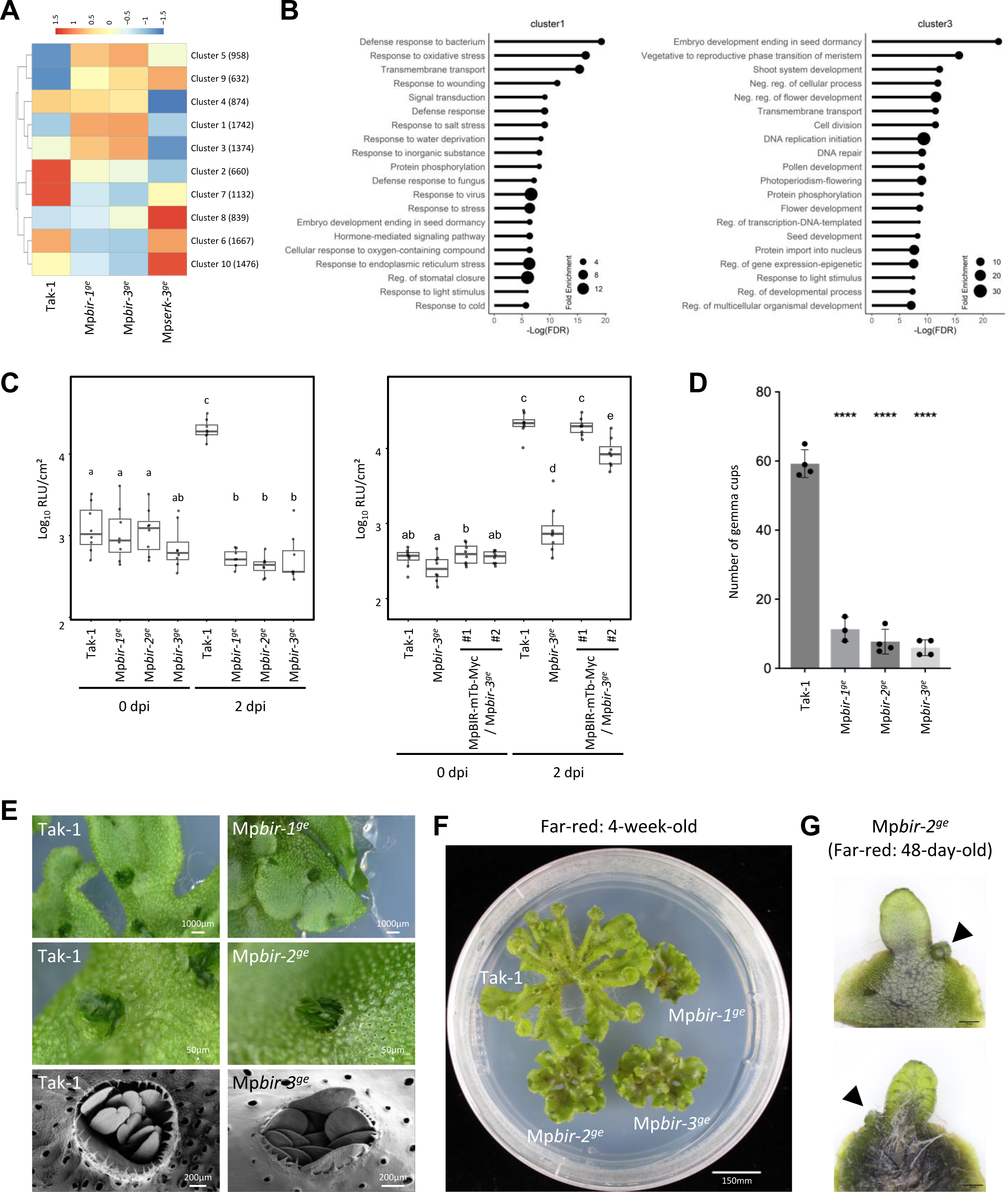
MpBIR negatively regulates immunity and is required for gemma cup and gametangiophore development. (A) Clusters of DEGs. Significantly differentially expressed genes with over ±1 log_2_ fold changes (false discovery rate [FDR]-adjusted p < 0.05) were grouped based on K-means clustering. Log_2_ read count of genes was normalized into the range of ±1.5. Number of genes in each cluster are shown in parentheses next to cluster IDs. See Data S1F. (B) Enriched GO terms in cluster 1 and cluster 3 shown in (A). See Data S1G and S1H. Quantification of bacterial growth in the central region of 14-day-old thalli. Plants were inoculated with the bioluminescent *Pto* DC3000-lux. Boxes show upper and lower quartiles of the value, and lines in boxes represent the medians (n = 8). Statistical analysis was performed using Student’s t test with p-values adjusted by the BH method. Statistically significant differences are indicated by different letters (p < 0.05). (C) Statistical analysis of the amount of gemma cups in Mp*bir-1^ge^*, Mp*bir-2^ge^*, Mp*bir-3^ge^,* and Tak-1. All plants were 29 days old. Significant differences calculated by one-way ANOVA are indicated with asterisks. Error bars, standard deviation. ****, p-value < 0.0001. (D) Gemma cups in 4-week-old Mp*bir-2^ge^* and Tak-1. Thalli were grown from single gemma under constant white light. Black and white images were taken by SEM. (E) Gametangiophore induction in 4-week-old Mp*bir-1^ge^*, Mp*bir-2^ge^*, Mp*bir-3^ge^*, and Tak-1. Thalli were grown from single gemma under constant far-red and white light. (F) Gametangiophore induction in 48-day-old Mp*bir-2^ge^*. Thalli were grown from single gemma under constant far-red and white light.

To investigate whether MpBIR or the MpSERK1MpBIR module actually contribute to defense against pathogenic bacteria, we challenged the Mp*bir^ge^* mutants with bioluminescent *Pseudomonas syringae* pv. *tomato* DC3000 (*Pto*-lux), and bacterial growth was measured at two days post-inoculation (dpi).^18,19^ Beyond our expectations, *Pto*-lux barely proliferated on 14-day-old Mp*bir^ge^* thalli, which strongly supports the transcriptome data (Figure 3C). This hyper-resistance phenotype could be reverted to the wild-type Tak-1 level by expression of MpBIR fused to mTb and Myc at its C-terminus (MpBIR-mTb-Myc) under the Mp*EF1α* promoter (Figure 3C). To understand how or whether MpSERK contributes to PRR-dependent immune signaling, we still need to identify PRRs that form complexes with MpSERK and their respective ligands. Nevertheless, our results indicate a role for the MpSERK1MpBIR module in *M. polymorpha* immunity.

Mp*bir^ge^* mutants also displayed defects in vegetative and sexual reproduction. Gemma cup formation was significantly reduced in all three Mp*bir^ge^* mutants (Figure 3D), and the mutants failed to establish the mature funnel-shaped cup (Figure 3E). In Mp*bir^ge^* mutants, far-red light irradiation triggered the induction of gametangiophore formation. However, gametangiophore development was disturbed at the early stage (Figures 3F and 3G). These results imply that MpBIR-dependent MpSERK regulation is required for gemma cup and gametangiophore development but not for induction of these processes.

### Phosphorylation status of various protein kinases is affected in Mpserk

To gain insight into phosphorylation events that are regulated by the MpSERK1MpBIR module, we profiled the phosphoproteomes of Tak-1, Mp*serk-3^ge^*, Mp*bir-1^ge^*, and Mp*bir-3^ge^*(Figure 4; Data S1I–S1M). Comparing the abundances of the identified phosphopeptides in Mp*serk-3^ge^* and Tak1, we found that more phosphopeptides were downregulated than upregulated in the mutant (Figures 4B and 4E; Data S1K–S1M). Meanwhile, an opposite pattern was observed in the Mp*bir^ge^* mutants (Figures 4B and 4E; Data S1K–S1M). These observations chime well with the assumption that MpSERK functions as a protein kinase and MpBIR constrains MpSERK activity.

**Figure 4.**
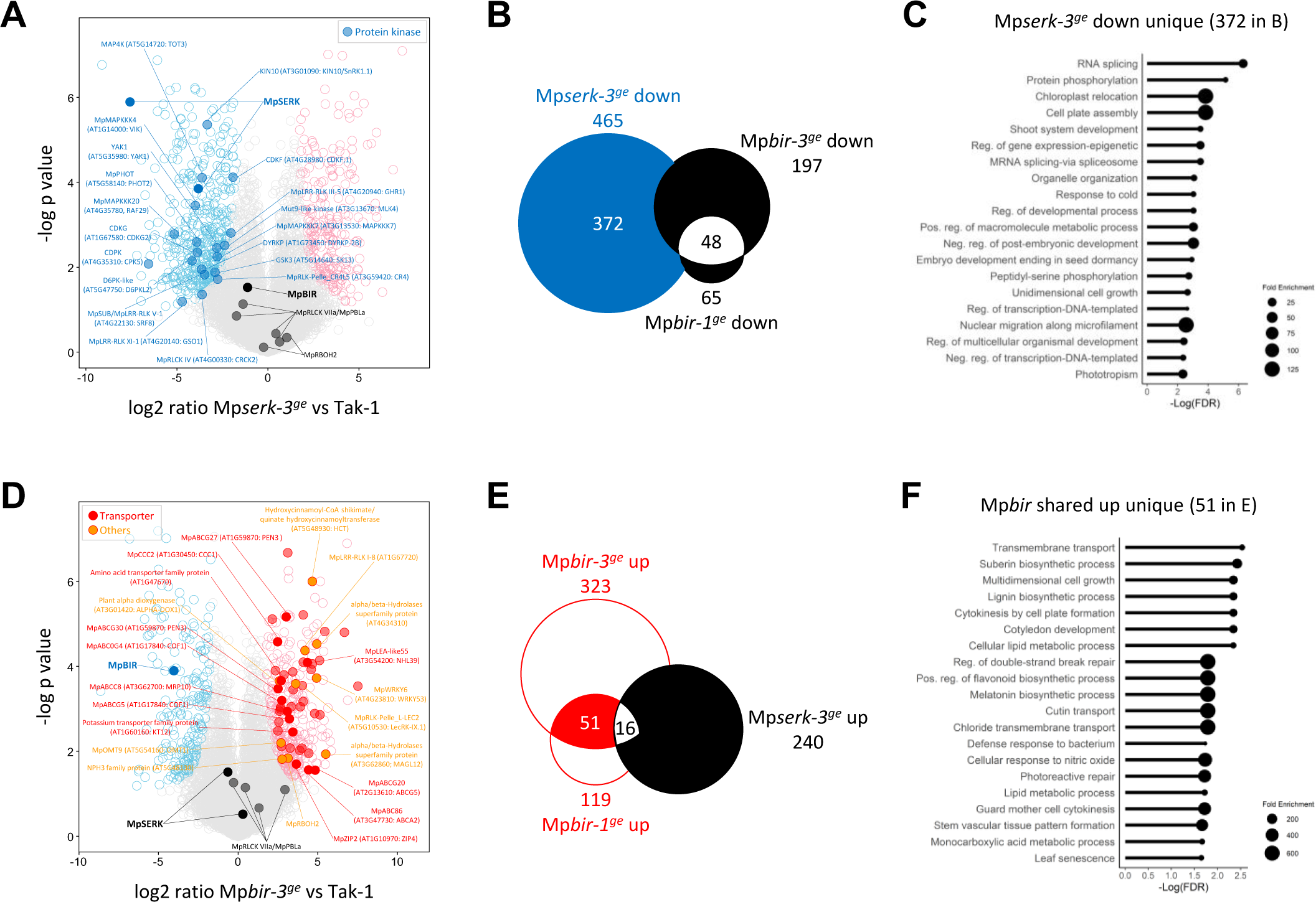
Phosphoproteome landscapes of Tak-1, Mpserk^ge^, and Mpbir^ge^. (A) A volcano plot showing differential abundance of phosphopeptides between Tak1 and Mp*serk-3^ge^*. Each dot represents a single unique phosphopeptide. Significantly upregulated and downregulated phosphopeptides in Mp*serk-3^ge^* compared to Tak-1 are colored red and blue, respectively (FDR = 0.05, *S0* = 1). Among significantly downregulated phosphopeptides, peptides derived from putative protein kinases are highlighted as filled circles with blue. Closest homologs in *A. thaliana* are shown in parentheses. See Data S1M. (B) Overlaps of the downregulated phosphopeptides in Mp*serk-3^ge^*, Mp*bir-1^ge^*, and Mp*bir-3^ge^* compared to Tak-1. Among 465 phosphopeptides downregulated in Mp*serk-3^ge^*, 372 phosphopeptides were specifically downregulated in Mp*serk-3^ge^* but not in Mp*bir^ge^* mutants. (C) GO enrichment analysis using 372 phosphopeptides identified in (B). See Data S1N and S1O. (D) A volcano plot showing differential abundance of phosphopeptides between Tak1 and Mp*bir-3^ge^*. Each dot represents a single unique phosphopeptide. Significantly upregulated and downregulated phosphopeptides in Mp*bir-3^ge^* compared to Tak-1 are colored red and blue, respectively (FDR = 0.05, *S0* = 1). Phosphopeptides upregulated both in Mp*bir-1^ge^* and Mp*bir-3^ge^* but not in Mp*serk-3^ge^* are shown as filled circles, except in blue and black (51 phosphopeptides identified in (E)). Among significantly upregulated phosphopeptides, peptides derived from putative transporters are highlighted as filled circles with red. Closest homologs in *A. thaliana* are shown in parentheses. See Data S1L. (E) Overlaps of the upregulated phosphopeptides in Mp*serk-3^ge^*, Mp*bir-1^ge^*, and Mp*bir-3^ge^*compared to Tak-1. Among 323 phosphopeptides upregulated in Mp*bir-3^ge^*, 51 phosphopeptides were also upregulated in Mp*bir-1^ge^* but not in Mp*serk-3^ge^*. (F) GO enrichment analysis using 51 phosphopeptides identified in (E). See Data S1P and S1Q. See also Data SI–K.

Among the proteins whose phosphorylation levels were downregulated in Mp*serk-3^ge^* but not differentially regulated in the Mp*bir^ge^* mutants, we found a number of protein kinases including LRR-RLK, receptor-like cytoplasmic kinase (RLCK), and MAP kinase (MAPK) cascade components (Figures 4A–4C; Data S1M–S1O). This was based on the assumption that phosphopeptide abundance reflects protein phosphorylation level. MpSUB/MpLRR-RLK V-1 (Mp4g21470), MpRLCK IV (Mp3g01680), MpPHOT (Mp5g03810), MAP4K (Mp8g10870), MpMAPKKK7 (Mp3g17970), and MpMAPKKK20 (Mp8g00050) were also identified as potential interactors of MpSERK (Figure 2A, 2B, and 4A; Data S1A and S1B), and therefore these kinases could be strong candidates that might be directly targeted by MpSERK. Other than protein kinases, phosphorylation levels of transcription factors (i.e., MpTRIHELIX1 (Mp1g16790), MpGRAS1 (Mp1g20490), MpWRKY1 (Mp3g00010), Mp1R-MYB20 (Mp4g00040)), RNA processing factor (i.e., MpSE (Mp1g23090)), auxin transporter (i.e., MpPIN1 (Mp3g21660)), calcium channel (i.e., MpOSCA1.2 (Mp7g04480)), and cell cycle regulator (i.e., MpRBR (Mp8g18830)) were down regulated in Mp*serk-3^ge^* (Data S1M). These factors may function downstream of MpSERK-dependent phosphorylation pathways for growth and development.

In the Mp*bir^ge^* mutants, our attention was caught by the upregulation of the phosphorylation levels of a number of transporters, including AtPEN3 homologs (i.e., MpABCG27 (Mp7g16260), and MpABCG30 (Mp2g21800)) (Figures 4D–4F; Data S1K, S1L, S1P, and S1Q). It is possible that the activities of these various transporters are precisely controlled by MpSERK, whose activity needs to be tightly repressed by MpBIR under non-stimulated or non-stressed conditions. Phosphorylation levels of proteins involved in extracellular matrix organization were also upregulated in the mutants (Figure 4D–4F; Data S1K, S1L, S1P, and S1Q). The hyper-resistance of the Mp*bir^ge^* mutants to *Pto*-lux can be explained by changes in extracellular physico-chemical properties (Figure 3C). Besides, increased phosphorylation of a WRKY transcription factor (i.e., MpWRKY6 (Mp1g08960)), LRR-RLK (i.e., MpLRR-RLK I-8 (Mp2g16600)), and lectin receptor-like kinase (LecRK) (i.e., MpRLK-Pelle_L-LEC2 (Mp1g13200)) was observed (Figures 4D–4F; Data S1K and S1L). Given that LecRKs positively regulate immunity in flowering plants, it is attractive to hypothesize that MpRLK-Pelle_L-LEC2 positively regulates immunity in *M. polymorpha* in conjunction with MpSERK.^23–25^

MpRBOH2 (synonymous with MpRBOH1 in Chu *et al*.^26^ and MpRBOHB in Hashimoto *et al*.^27^) plays roles in development and the chitin-induced reactive oxygen species (ROS) burst in *M. polymorpha*. Chitin-induced responses are mediated through LysM-receptor-like kinase (LysM-RLK: LYK) MpLYK1 and LYK-related (LYR) MpLYR.^18^ MpRLCK VIIa/MpPBLa phosphorylates MpRBOH2 *in vitro* and is indispensable for the chitin-induced ROS burst.^26^ In the Mp*bir^ge^* mutants, the phosphorylation level of MpRBOH2 was increased (Figure 4D). We also observed a trend towards increased phosphorylation of MpRLCK VIIa in the Mp*bir^ge^*mutants (Figure 4D). These results may imply that MpRLCK VIIa functions downstream of both LysM-RLK and LRR-RLK.

## Conclusion

We have shown here that in the liverwort *M. polymorpha* SERK and BIR function together as a module that plays crucial roles both in development and immunity. Antagonistic transcriptome profiles of Mp*serk^ge^*and Mp*bir^ge^* mutants support our hypothesis that MpBIR suppresses MpSERK-mediated pathways. These findings demonstrate that physiological and molecular functions of the SERK–BIR module in development and immunity are highly conserved across land plants. This suggests that the SERK–BIR module evolved in the MRCA of land plants. The next challenge will be to identify receptor kinases and their ligands which function together with MpSERK and regulate development or immunity. We expect that the transcriptome, interactome, and phosphoproteome data reported in this study will facilitate further dissection of SERK-mediated pathways and their evolution.

## Resource availability

### Lead contact

Requests for resources and further information should be directed towards Hirofumi Nakagami.

### Materials availability

Plasmids and plant materials generated in this study are all available upon request. Please note that the distribution of transgenic plants will be governed by material transfer agreements (MTAs) and will be dependent on appropriate import permits acquired by the receiver.

## Experimental model and subject details

### Plant materials and growth condition

*Marchantia polymorpha* accession Tak-1 was used as a wild-type throughout this study. For cultivation, gemmae were grown on half-strength Gamborg’s B5 (GB5) basal media containing 1% agar at 22 °C under continuous white light (60 to 70 μmol m^-2^ s^-1^). For Mp*serk^ge^* mutants, small fragments that were cut from the apical region were used as starting materials instead of gemmae. For gametangiophore induction, plants were grown under continuous white light (60–70 μmol m^-2^ s^-1^) supplemented with far-red light (60–65 μmol m^-2^ s^-1^).

## Method details

### Phylogenetic analysis

Protein sequences were retrieved from the following databases: Phytozome (https://phytozome-next.jgi.doe.gov/), MarpolBase (https://marchantia.info) and TAIR (http://www.arabidopsis.org/). Alignment was performed on the amino acid sequences of the kinase domain using CLUSTALW (https://www.genome.jp/tools-bin/clustalw). Phylogenetic analysis was performed on the alignment using MrBayes3.2.769. Two runs with four chains of Markov chain Monte Carlo (MCMC) iterations were performed for 300,000 generations for SERK or 600,000 generations for BIR, keeping one tree every 100 generations. The first 25% of generations were discarded as burn-in and the remaining trees were used to calculate a 50% majority-rule tree. The standard deviation for the two MCMC iteration runs was below 0.01, suggesting that it was sufficient for the convergence of the two runs. Convergence was assessed by visual inspection of the plot of the log likelihood scores of the two runs calculated by MrBayes.

### DNA/RNA extraction and cDNA synthesis

Fresh thalli were frozen and ground using a MM 400 mixer mill (Retsch, Germany). DNA and RNA were extracted using the DNeasy Plant Mini Kit and RNeasy Plant Mini Kit (QIAGEN, Netherlands), respectively. cDNA synthesis was performed using SuperScript IV Reverse Transcriptase (Invitrogen, USA).

### Plasmid constructions and transformation

To construct the vectors for generating Mp*serk^ge^* and Mp*bir^ge^* mutants, annealed oligos for Mp*SERK*- and Mp*BIR-*targeting gRNAs were ligated into *Bsa*I-digested pMpGE_En03^28^ using a T4 DNA ligase (NEB, UK). Mp*SERK*- and Mp*BIR-*targeting gRNAs were then subcloned into the binary vectors pMpGE011 and pMpGE010, respectively. The plasmid containing Mp*SERK*-targeting gRNA was introduced into Tak-1, Mp*bir-3^ge^*, and Mp*bir-1^ge^* using the *Agrobacterium*-mediated transformation method^29^ to generate Mp*serk^ge^*, Mp*serk-4^ge^*/Mp*bir-3^ge^*, and Mp*serk-5^ge^*/Mp*bir-1^ge^*, respectively. The plasmid containing Mp*BIR*-targeting gRNA was introduced into Tak-1 and Mp*serk-1^ge^* to generate Mp*bir^ge^* and Mp*bir^ge^*/Mp*serk-1^ge^*, respectively. Screening for CRISPR/Cas9-mediated targeted mutagenesis was performed by genomic PCR as described previously.^30^ The 5 kb putative promoter fragments upstream of the translation initiation codon of Mp*SERK* and Mp*BIR* were cloned into the Gateway^TM^ pENTR4 dual-selection vector (Thermo Fisher Scientific, USA) using an In-Fusion HD cloning kit (Takara, Japan), followed by subcloning into the pMpGWB304 binary vector for construction of *_pro_*Mp*SERK:GUS* using LR clonase II enzyme mix (Thermo Fisher Scientific, USA).^31^ The Mp*BIR* promoter was cloned into Gateway^TM^ pDNOR207 vector (Thermo Fisher Scientific, USA) using the NEBuilder® HiFi DNA Assembly Cloning Kit (NEB, UK). The coding sequences of Mp*SERK* and Mp*SERK^Y418F^* were synthesized (GeneArt Gene Synthesis; Thermo Fisher Scientific, USA). The gRNA target site and PAM sequence of Mp*SERK* were mutated, so as not to be targeted by CRISPR/Cas9 in Mp*serk^ge^*. Synthesized coding sequences of Mp*SERK* and Mp*SERK^Y418F^* were cloned into pENTR4-*_pro_*Mp*SERK*, and then they were subcloned into binary vector pMKMM1 (Figure S3) for construction of *_pro_*Mp*SERK:*Mp*SERK-miniTurbo-Myc* and *_pro_*Mp*SERK:*Mp*SERK^Y418F^-miniTurbo-Myc*. The resulting plasmids were introduced into Mp*serk-3^ge^*. The coding sequence of Mp*BIR* was amplified from cDNA prepared from Tak-1 using KOD Plus Neo (Toyobo, Japan). Mp*BIR* coding sequence was cloned into pDNOR207 and pDNOR207-_pro_Mp*BIR* vectors, followed by subcloning into binary vectors pMKMM2 and pMKMM1 (Figures S3) to construct *_pro_*Mp*EF1α:*Mp*BIR-miniTurbo-Myc* and *_pro_*Mp*BIR:*Mp*BIR-miniTurbo-Myc*, respectively. The *_pro_*Mp*EF1*α:Mp*BIR-miniTurbo-Myc* plasmid was introduced into Tak-1 and *_pro_*Mp*BIR:*Mp*BIR-miniTurbo-Myc* plasmid was introduced into Mp*bir-3^ge^*. To generate estradiol-inducible expression vectors for transient expression in *N. benthamiana* leaves, Mp*SERK* and Mp*BIR* were subcloned into binary vector pABind117 and pABind118, respectively^32^, to construct Mp*SERK:mCherry* and Mp*BIR:GFP* using LR clonase II enzyme mix (Thermo Fisher Scientific, USA). The used oligonucleotides are listed in Table S1.

### Cryo-scanning electron microscopy (Cryo-SEM)

Samples were mounted on copper sample holders, snap-frozen in liquid nitrogen and sublimated, sputtered with Gold/Palladium mixture (80% Gold/20% Palladium) using an Emitech K1250X cryo system, and then images were taken using a Zeiss Supra 40VP scanning electron microscope.

### GUS histochemical assay

Seven or 14-day-old thalli were submerged in GUS staining solution consisting of 0.5 mg/mL X-Gluc (5-bromo-4-chloro-3-indolyl-beta-D-glucuronic acid), 0.1% Triton X-100, 10 mM EDTA, 0.5 mM potassium ferricyanide, 0.5 mM potassium ferrocyanide in 100 mM sodium phosphate buffer (pH 7.0), followed by vacuum infiltration for 5–15 minutes. After overnight incubation in the dark at 37 °C, tissues were de-stained by incubation in 70% ethanol with gentle shaking for a minimum of 1 hour before observation. For sectioning, GUS-stained samples were embedded into 5% agarose. Embedded samples were then sectioned into 150 μm-thick sections using VT1000 S vibratome (Leica, Germany).

### Immunoblotting

Proteins were separated by sodium dodecyl sulphate (SDS)-polyacrylamide-gel electrophoresis (PAGE) and blotted onto polyvinylidene fluoride (PVDF) membranes (1704272; Bio-Rad, USA) using a Trans-Blot Turbo (Bio-Rad, USA) transfer system. The membrane was probed with anti-Myc-tag mouse monoclonal antibody (9B11; Cell Signaling Technology, USA) overnight at 4 °C and then with horseradish peroxidase (HRP)-conjugated anti-mouse immunoglobulin G (IgG) antibody (7076S; Cell Signaling Technology, USA) for 1 hour at room temperature. Proteins were visualized on the membrane using a luminol-based chemiluminescent substrate that is oxidized by HRP in the presence of peroxide (34577; Thermo Fisher Scientific, USA). The membranes were then stained with Ponceau S for 10 minutes at room temperature and then rinsed with distilled water.

### miniTurbo-based interactomics

Interactome analysis was carried out as described before.^20,33^ Briefly, 14-day-old thalli were collected, vacuum-infiltrated with 700 μM biotin solution, and incubated overnight at room temperature in biotin solution with gentle shaking. After incubation, thalli were washed with Milli-Q water, drained on filter paper, and snap-frozen in liquid nitrogen. Plants were ground into tissue powder and the total protein was extracted. Then, 500 μg of total protein were used for biotin depletion by methanol–chloroform precipitation. Biotinylated proteins were pulled-down using Streptavidin Mag Sepharose^TM^ (Cytiva 28-9857-99; GE Healthcare) and then submitted to on-bead digestion. The beads were resuspended in 25 µL digestion buffer 1 (50 mM Tris pH 7.5, 2 M urea, 1 mM dithiothreitol (DTT), 5 µg/ml Trypsin) and incubated in a thermomixer at 32 °C with agitation at 400 rpm for 30 minutes. The supernatant was transferred to a fresh tube. The beads were then treated with 50 µl digestion buffer 2 (50 mM Tris pH 7.5, 2 M Urea, 5 mM chloroacetamide (CAA)). Obtained supernatants were combined and the total digest was incubated overnight in a thermomixer at 32 °C with agitation at 400 rpm. After acidification with 10% trifluoroacetic acid (TFA), samples were desalted with C18 Empore disk membranes according to the StageTip protocol.^34^ The eluted peptides were dried and then dissolved in 101µl buffer A (2% acetonitrile (ACN), 0.1% TFA) and measured without dilution.

Samples were analyzed using an EASY-nLC 1200 (Thermo Fisher Scientific, USA) coupled to a Q Exactive Plus mass spectrometer (Thermo Fisher Scientific, USA). Peptides were separated on 16-cm frit-less silica emitters (75 μm inner diameter; New Objective, USA), packed in-house with reversed-phase ReproSil-Pur C18 AQ 1.9 µm resin (Dr. Maisch, Germany). Peptides were loaded on the column and eluted for 60 minutes using a segmented linear gradient of 5% to 95% solvent B (0 min: 5%B; 0–5 min → 5%B; 5–25 min → 15%B; 25–50 min → 35%B; 50–55 min → 95%B; 55–60 min → 95%B) (solvent A 0% ACN, 0.1% FA; solvent B 80% ACN, 0.1%FA) at a flow rate of 300 nl/min. Mass spectra were acquired in data-dependent acquisition mode with a TOP12 method. MS spectra were acquired in the Orbitrap analyzer with a mass range of 300–1,500 m/z at a resolution of 70,000 FWHM and a target value of 3 × 10^6^ ions. Precursors were selected with an isolation window of 1.3 m/z. HCD fragmentation was performed at a normalized collision energy of 25. MS/MS spectra were acquired with a target value of 5 x 10^5^ ions at a resolution of 17,500 FWHM, a maximum injection time of 85 ms and a fixed first mass of m/z 100. Peptides with a charge of 1, greater than 6, or with unassigned charge state were excluded from fragmentation for MS^2^. Dynamic exclusion for 20 seconds prevented repeated selection of precursors.

Raw data were processed using MaxQuant software (version 1.6.3.4, http://www.maxquant.org/)^35^ with label-free quantification (LFQ) and iBAQ enabled^36^; normalization was skipped for the LFQ quantification. MS/MS spectra were searched by the Andromeda search engine against a combined database containing the sequences from *M. polymorpha* (MpTak1v5.1_r1_primary_transcripts_proteinV3; https://marchantia.info/) and sequences of 248 common contaminant proteins and decoy sequences and the sequence of the miniTurbo. Trypsin specificity was required and a maximum of two missed cleavages allowed. Minimal peptide length was set to seven amino acids. Carbamidomethylation of cysteine residues was set as fixed, oxidation of methionine and protein N-terminal acetylation as variable modifications. Peptide-spectrum-matches and proteins were retained if they were below a false discovery rate of 1%. The non-normalized MaxLFQ values of every two-genotype combination (five replicates per condition) were pre-processed in Perseus (version 1.5.8.5, http://www.maxquant.org/) and submitted for normalization analysis using the Normalyzer tool (http://normalyzer.immunoprot.lth.se/).^37^ The output was analyzed for outliers and one replicate per condition was removed in the subsequent data analysis. The final data analysis was carried out in MaxQuant as described above on the reduced raw dataset; each two-genotype combination was searched independently. Statistical analysis of the MaxLFQ values was carried out using Perseus. Quantified proteins were filtered for reverse hits and hits “identified by site” and MaxLFQ values were log_2_ transformed and the data was normalized by subtraction of the median per column. After grouping samples by condition only those proteins were retained for the subsequent analysis that had three valid values in one of the conditions. Two-sample *t*-tests were performed using a permutation-based FDR of 5%. The output was exported to Excel for further processing. Alternatively, data was filtered for three valid values in one of the conditions and missing values were imputed from a normal distribution (1.8 downshift, separately for each column). Volcano plots were generated in Perseus using an FDR of 5% and an *S0* = 1. The Perseus output was exported and further processed using Excel and RStudio.

### Transient expression in *Nicotiana benthamiana*

*Agrobacterium* GV3101 strains transformed with the desired vectors were cultured on Luria-Bertani (LB) plate (1% agar) containing the respective antibiotics for two days at 28 °C. A single colony was selected and inoculated into 5 ml of LB medium with the appropriate antibiotics, then incubated overnight at 28 °C with shaking. To prepare a fresh liquid culture, 1 ml of the overnight culture was added to 4 ml of LB medium containing antibiotics and incubated with shaking at 28 °C for 4 hours. The bacteria were then collected by centrifugation and resuspended in 5 ml of infiltration solution (5% sucrose, 0.01% Silwet® L-77, 450 μM acetosyringone, a spatula tip of glucose). The bacterial suspension was kept on ice before infiltration. The optical density (OD_600_) was measured using a spectrophotometer and adjusted to an OD_600_ of 0.4 per strain using fresh infiltration solution. *Nicotiana benthamiana* leaves were infiltrated with the suspension using a needleless syringe, and the infiltrated areas were marked with a permanent marker. The inflated plants were kept under continuous white LED (50–60 µmol photons m^-2^s^-1^) at 22 °C for at least 48 hours. Then, the abaxial side of the infilled leaves were painted with an induction solution (20 μM β-estradiol, 0.1% Tween 20) to induce protein expression 24 hours before observation.

### FLIM-FRET

*Nicotiana benthamiana* leaf samples expressing either only MpBIR-GFP, as donor in absence of MpSERK-mCherry as acceptor, or in combination with MpSERK-mCherry, were mounted on microscope slides in water, covered with a high-precision cover glass and immediately used for analysis of fluorescence life times. For this, a Leica SP8 FALCON-DIVE confocal system, equipped with an InSight X3 pulsed laser from Spectra Physics with a fixed laser line of 1,045 and a line tunable from 680 to 1,300 nm, was used in combination with either a 40x/1.25 NA GLYC or 40x/1.10 W immersion objective. For imaging and FLIM experiments, GFP was excited with 930 nm and the emission window from 490 to 550 nm was recorded with the RLD detector. To observe FRET between GFP as donor and mCherry as acceptor, only the donor fluorescence was recorded for lifetime imaging. Images with a frame size of 512 by 512 pixels were acquired until a level of 1,000 photons was reached for the maximum pixel value. Mean τ intensity weighted lifetimes (ns) were averaged across multiple regions of interest, containing two neighboring cells.

### RNA-Seq and data analysis

Total RNA was isolated from 14-day-old thalli grown on the agar plates using RNeasy Plant mini Kits (QIAGEN, Netherlands). Library preparation and sequencing were conducted by Novogene, UK (https://www.novogene.com/eu-en/) using the Illumina NovaSeq 6000 platform. The *M. polymorpha* genome version MpTak_v6.1r1 (https://marchantia.info/) was used for mapping and counting transcripts per gene in STAR aligner.^38^ Genes with less than the average of 10 read counts were excluded, and DESeq2 was used for raw count normalization and differentially expressed gene (DEG) analyses.^39^ Statistically significant DEGs (adjusted p-value < 0.05) were selected for further analyses.

### GO analysis

Corresponding Arabidopsis gene IDs were annotated to *M. polymorpha* using DIAMOND with an e-value cutoff of 0.0001.^40^ GO enrichment analyses were performed on selected DEGs and proteins in ShinyGO using Arabidopsis DB with FDR cutoff of 0.05. The top 20 GO biological processes are shown.^41^

### Bioluminescence-based bacteria quantification

Bacterial quantification in infected thalli was carried out as described before.^19^ Briefly, *M. polymorpha* were grown on autoclaved cellophane discs on half-strength GB5 media for 2 weeks. In the meantime, *Pto*-lux was cultivated in King’s B medium containing 30 μg/ml rifampicin to achieve an OD_600_ of 1.0. The saturated bacterial culture was subsequently washed and resuspended in Milli-Q water to prepare a bacterial suspension with an 0.01 of OD_600_. Next, 2-week-old thalli were submerged in the bacterial suspension followed by vacuum for 5 minutes and incubated for 0 to 2 days on humid filter papers. After incubation, thallus discs (5 mm diameter) were punched from the central region using a sterile biopsy punch (pfm medical, Germany) and transferred to a 96-well plate. The bioluminescence was measured in the FLUOstar Omega plate reader (BMG Labtech, Germany).

### Phosphoproteomics

Phosphoproteome analysis was carried out as described before with minor modifications.^18,42^ Fourteen-day-old thalli were snap-frozen in liquid nitrogen and were disrupted using a MM 400 mixer mill (Retsch, Germany). Sample preparation was performed as described previously with minor modifications.^43^ Samples were analyzed using an Ultimate 3000 RSLC nano (Thermo Fisher Scientific, USA) coupled to an Orbitrap Exploris 480 mass spectrometer equipped with a FAIMS Pro interface for Field asymmetric ion mobility separation (Thermo Fisher Scientific, USA). Peptides were pre-concentrated on an Acclaim PepMap 100 pre-column (75 µM x 2 cm, C18, 3 µM, 100 Å; Thermo Fisher Scientific, USA) using the loading pump and buffer A (water, 0.1% TFA) with a flow of 7 µl/min for 5 minutes. Peptides were separated on 16 cm frit-less silica emitters (75 μm inner diameter, New Objective, USA), packed in-house with reversed-phase ReproSil-Pur C18 AQ 1.9 µm resin (Dr. Maisch, Germany). Peptides were loaded on the column and eluted for 130 minutes using a segmented linear gradient of 5% to 95% solvent B (0 min: 5% B; 0–5 min → 5% B; 5–65 min → 20% B; 65–90 min → 35% B; 90–100 min → 55% B; 100–105 min → 95% B, 105–115 min → 95% B, 115–115.1 min → 5% B, 115.1–130 min → 5% B) (solvent A 0% ACN, 0.1% FA; solvent B 80% ACN, 0.1%FA) at a flow rate of 300 nl/min. Mass spectra were acquired in data-dependent acquisition mode with a TOP_S method using a cycle time of 2 seconds. For field asymmetric ion mobility separation (FAIMS) two compensation voltages (−45 and −65) were applied, the cycle time for the CV-45 experiment was set to 1.2 seconds and for the CV-65 experiment to 0.8 seconds. MS spectra were acquired in the Orbitrap analyzer with a mass range of 320–1,200 m/z at a resolution of 60,000 FWHM and a normalized AGC target of 300%. Precursors were filtered using the MIPS option (MIPS mode = peptide), the intensity threshold was set to 5,000, and precursors were selected with an isolation window of 1.6 m/z. HCD fragmentation was performed at a normalized collision energy of 30%. MS/MS spectra were acquired with a target value of 75% ions at a resolution of 15,000 FWHM, at an injection time of 120 ms and a fixed first mass of m/z 120. Peptides with a charge of +1, greater than 6, or with unassigned charge state were excluded from fragmentation for MS^2^.

Raw data were processed using MaxQuant software (version 1.6.3.4, http://www.maxquant.org/)^35^ with label-free quantification (LFQ) and iBAQ enabled.^36^ MS/MS spectra were scanned by the Andromeda search engine against a combined database containing the sequences from *M. polymorpha* (MpTak1v6.1_r2.protein.fasta; https://marchantia.info/), sequences of 248 common contaminant proteins, and decoy sequences. Trypsin specificity was required and a maximum of two missed cleavages allowed. Minimal peptide length was set to seven amino acids. Carbamidomethylation of cysteine residues was set as fixed, phosphorylation of serine, threonine and tyrosine, oxidation of methionine and protein N-terminal acetylation as variable modifications. The match between runs option was enabled. Peptide-spectrum-matches and proteins were retained if they were below a false discovery rate of 1% in both cases. Statistical analysis of the intensity values obtained for the phospho-modified peptides (“modificationSpecificPeptides.txt” output file) was carried out using Perseus (version 1.5.8.5, http://www.maxquant.org/). Intensities were filtered for reverse and contaminant hits and the data was filtered to retain only phospho-modified peptides. Next, intensity values were log_2_-transformed. After grouping samples by condition, only those sites were retained for the subsequent analysis that had three valid values in one of the conditions. Two-sample t-tests were performed using a permutation-based FDR of 0.05. Alternatively, data were filtered for four valid values and were median-normalized. Missing values were imputed from a normal distribution, using the default settings in Perseus (1.8 downshift, separately for each column). Volcano plots were generated using FDR = 0.05 and *S0* = 1. The Perseus output was exported and further processed using Excel and RStudio.

### Quantification and statistical analysis

Excel, R (version 4.2.3), RStudio (version 2024.04.1), and Prism 9.0 (GraphPad Software) were used for statistical analysis and figure preparation. Detailed statistical information, including sample sizes and error bars are provided in the corresponding figure legends. Bacterial growth was analyzed using Student’s t-test, with p-values adjusted by the Benjamini and Hochberg (BH) method. Statistically significant differences were defined as p < 0.05. Quantifications of GFP fluorescence lifetime and the number of gemma cups were analyzed using one-way ANOVA followed by Dunnett’s test, comparing each condition to the control, using Prism 9.0.

## Data availability

Sequencing raw reads used in transcriptomic analyses in this study have been deposited under the accession BioProject PRJNA1128533. The MS proteomics data have been deposited in the ProteomeXchange Consortium via the PRIDE partner repository with the dataset identifiers PXD053428 and PXD053600.

## Acknowledgments

We thank Ton Timmers (MPIPZ, Germany), Tonni Grube Andersen (MPIPZ, Germany), and Gabriel Xicoténcatl García Ramírez (MPIPZ, Germany) for their technical support with the microscopic analyses. We thank Neysan Donnelly (MPIPZ, Germany) for editing the manuscript. This project was supported by the Max Planck Society and was conducted in the framework of MAdLand (http://madland.science, Deutsche Forschungsgemeinschaft [DFG] Priority Programme 2237). H.N. is grateful for funding by the DFG (NA 946/1-1 and NA 946/1-2). Y.H. was supported by JSPS KAKENHI grant number JP22H02676. Y.Y. is grateful to the China Scholarship Council (CSC) for PhD studentship funding.

## Author contributions

Y.Y., A.I.C.-D., and H.N. designed the research. Y.Y., J.M., M.A.M., H.-W.J., K.M., C.H., and Y.H. generated materials. Y.H. performed phylogenetic analysis. Y.Y, J.M., M.A.M., Y.H., and H.N. performed microscopic analysis. R.F. performed Cryo-SEM. Y.Y. and Y.H. performed GUS histochemical assay. Y.Y., A.H., S.C.S., Y.Y., and H.N. performed interactome analysis. Y.Y., H.-W.J., and H.N. performed transcriptomic analysis. Y.Y. and J.M. performed *Pto* DC3000 infection assay. Y.Y., A.H., S.C.S., Y.Y., and H.N. performed phosphoproteome analysis. Y.Y. performed all other experiments. Y.Y., H.-W.J., and H.N. wrote the manuscript. All authors corrected the manuscript.

## Supplemental information

**Data S1. Information on interactomics results, RNA-seq data analysis results, and phosphoproteomics results used in this study, related to Figures 2, 3, 4, and S1**

(A) Interactome data, related to Figure 2A. Comparison between MpSERK and Tak-1 for drawing volcano-plot.

(B) Interactome data, related to Figure 2B. Comparison between MpSERK^Y418F^ and Tak-1 for drawing volcano-plot.

(C) Interactome data, related to Figure 2C. Comparison between MpSERK and MpSERK^Y418F^ for drawing volcano-plot.

(D) Interactome data, related to Figure S1A. Comparison between MpSERK and MpSYP13B for drawing volcano-plot.

(E) Interactome data, related to Figure S1B. Comparison between MpSERKY^418F^ and MpSYP13B for drawing volcano-plot.

(F) RNA-seq data, related to Figure 3A.

(G) GO enrichment analysis of cluster 3, related to Figure 3B.

(H) GO enrichment analysis of cluster 1, related to Figure 3B.

(I) Phosphoproteome data, related to Figure 4. Data before imputation.

(J) Phosphoproteome data, related to Figure 4. Histogram.

(K) Phosphoproteome data, related to Figure 4. Comparison between Mp*bir-1^ge^* and Tak-1 for drawing volcano-plot.

(L) Phosphoproteome data, related to Figure 4. Comparison between Mp*bir-3^ge^* and Tak-1 for drawing volcano-plot.

(M) Phosphoproteome data, related to Figure 4. Comparison between Mp*serk-3^ge^* and Tak-1 for drawing volcano-plot.

(N) List of proteins used for GO enrichment analyses, related to Figure 4C.

(O) GO enrichment analysis, related to Figure 4C.

(P) List of proteins used for GO enrichment analyses, related to Figure 4F.

(Q) GO enrichment analysis, related to Figure 4F.

