## Supplemental Figures for "Conserved role of the SERK–BIR module in development and immunity across land plants"

### Figure S1

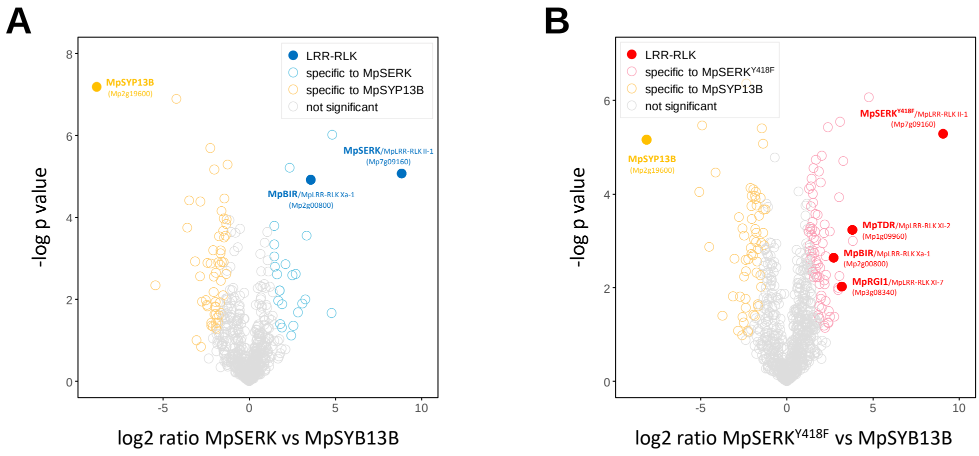


**Figure S1. Interactome analysis of MpSERK and MpSERK$^{Y418F}$ using MpSYP13B as a control, related to Figure 2**

(A) Proteins significantly enriched using MpSERK as a bait are indicated in blue circles. LRR-RLKs are highlighted as filled blue circles. (B) Proteins significantly enriched using MpSERK$^{Y418F}$ as a bait are indicated in red circles. LRR-RLKs are highlighted as filled red circles.

**Figure S2**

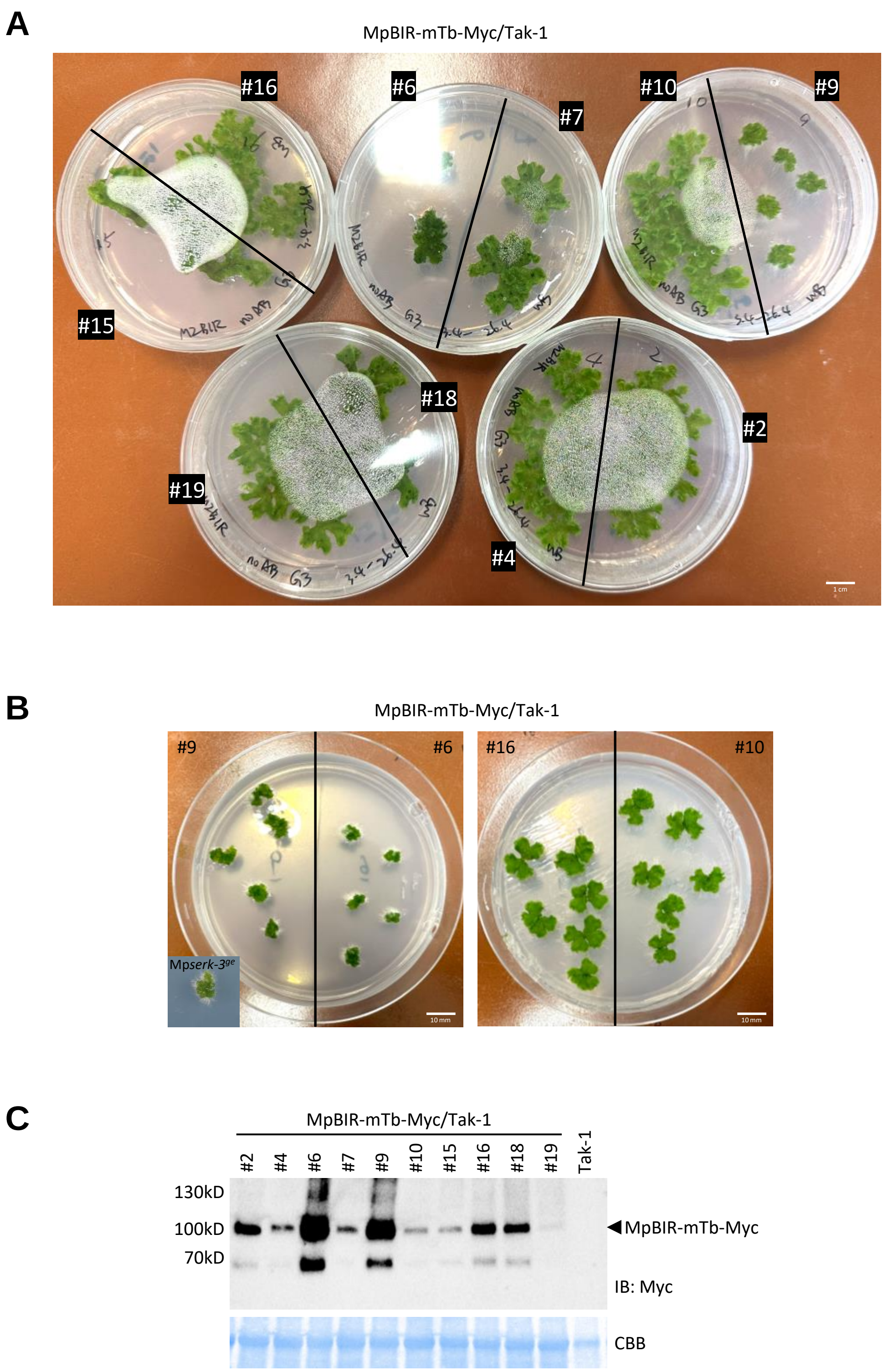


**Figure S2. MpBIR overexpressing plants, related to Figure 2**

(A) Isolated transgenic lines expressing Mp*BIR* under Mp*EF1α* promoter in Tak-1. Scale bar, 10 mm.

(B) Two-week-old $_{pro}$Mp*EF1α*:Mp*BIR-miniTurbo-Myc*/Tak-1 plants grown on agar plates. Thalli were grown from single gemma (line #10 and #16) or small fragments of thalli (line #6, #9, and Mp*serk-3*$^{ge}$). Scale bars, 10 mm.

(C) Protein expression levels of MpBIR-miniTurbo-Myc in the transgenic plants shown in (A and B). Myc-tagged proteins were detected using anti-Myc antibody and indicated by arrows. CBB-stained membrane is shown as a loading control.

### Figure S3

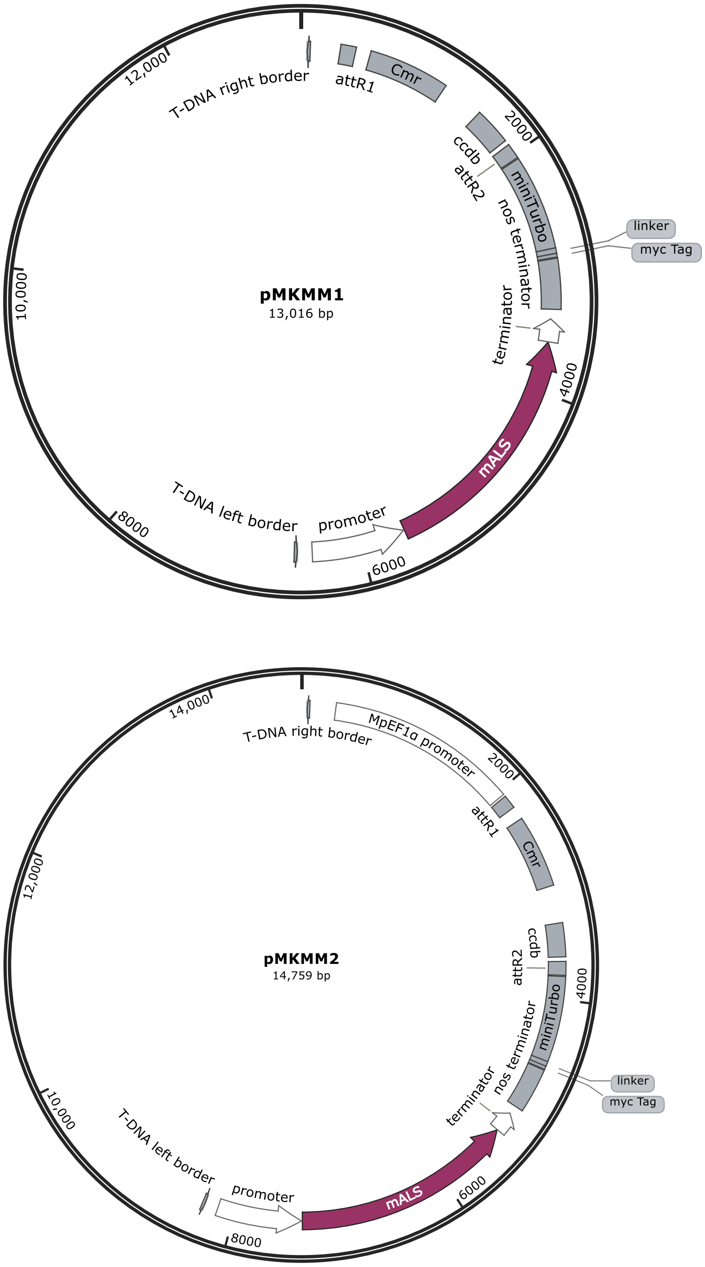


**Figure S3. Plasmid maps of pMKMM1 and pMKMM2, related to Method details**

| Name | Description | Sequence |
|---|---|---|
| MpSERK_gRNA_F | Construction of targeting vector for Mp*serk*$^{ge}$ generation | CTCGCAAGGGTCGTCTGGCTGA |
| MpSERK_gRNA_R | | AAACTCAGCCAGACGACCCTTG |
| MpBIR1_65_gRNA_F | Construction of targeting vector for Mp*bir*$^{ge}$ generation | CTCGAGACAAATCCAAGGAAGTAA |
| MpBIR1_65_gRNA_R | | AAACTTACTTCCTTGGATTTGTCT |
| pET4-pMpSERK-MpSERK_inF | MpSERK CDS amplification | AATCTCTGGAAAAGACTAGTCATGCAGCATCCTTGGTTCC |
| pET4-pMpSERK-MpSERK_inR | | GTACAAGAAAGCTGGGTCTCTTG |
| attB1-MpSERK | | GGGGACAAGTTTGTACAAAAAAGCAGGCTATGCAGCATCCTTGGTTCCT |
| attB2-MpSERK | | GGGGACCACTTTGTACAAGAAAGCTGGGTCTCTTGGGCCTGACAGTTCGA |
| MpSERKproF | MpSERK promoter amplification | GGGGGGATCCAAAATATTGCGGACTAGC |
| MpSERKproR | | GGGGGAATTCGACTAGTCTTTTCCAGAGATTCC |
| attB1-BIR | MpBIR CDS amplification | GGGGACAAGTTTGTACAAAAAAGCAGGCTATGTCTTTGGAAAACCCAGAGT |
| attB2-BIR | | GGGGACCACTTTGTACAAGAAAGCTGGGTCTGAGTTAGATACAATCAGCTCCT |
| pBIR-BIR-hfR | MpBIR promoter amplification | TTTGTACAAAAAAGCAGGCTAAACTCGTCTGATCTCACTCAACG |
| 207-pBIR-hfF | | TCTGGGTTTTCCAAAGACATCGCTCCCCGGCTTTGAATG |
| MpBIRseq-F | Genotyping Mp*bir*$^{ge}$ | ATGTCTTTGGAAAACCCAGAGTTG |
| MpSERK_seq.F2 | Genotyping Mp*serk*$^{ge}$ | GATGTCCCAGCTGAAGAAGA |

**Table S1. Oligonucleotides used in this study, related to Method details**
